# Superlets: time-frequency super-resolution using wavelet sets

**DOI:** 10.1101/583732

**Authors:** Vasile V. Moca, Adriana Nagy-Dăbâcan, Harald Bârzan, Raul C. Mureşan

**Affiliations:** Transylvanian Institute of Neuroscience, Department of Experimental and Theoretical Neuroscience, Pta. Timotei Cipariu 9/20, 400191 Cluj-Napoca, Romania; Technical University of Cluj-Napoca, Basis of Electronics Department, Str. G. Baritiu 26-28, 400027 Cluj-Napoca, Romania

**Keywords:** superlets, time-frequency analysis, neural oscillations, local-field potentials, electroencephalography, gamma bursts

## Abstract

Time-frequency analysis is ubiquitous in many fields of science. Due to the Heisenberg-Gabor uncertainty principle, a single measurement cannot estimate precisely the location of a finite oscillation in both time and frequency. Classical spectral estimators, like the short-time Fourier transform (STFT) or the continuous-wavelet transform (CWT) optimize either temporal or frequency resolution, or find a tradeoff that is suboptimal in both dimensions. Following concepts from optical super-resolution, we introduce a new spectral estimator enabling time-frequency super-resolution. Sets of wavelets with increasing bandwidth are combined geometrically in a *superlet* to maintain the good temporal resolution of wavelets and gain frequency resolution in the upper bands. *Superlets* outperform the STFT, CWT, and other super-resolution methods on synthetic data and brain signals recorded in humans and rodents, resolving time-frequency details with unprecedented precision. Importantly, *superlets* can reveal transient oscillation events that are hidden in the averaged time-frequency spectrum by other methods.

## Introduction

Time-series describing natural phenomena, such as sounds, earth movement, or brain activity, often express oscillation bursts, or “packets”, at various frequencies and with finite duration. In brain signals, these packets span a wide range of frequencies (e.g., 0.1-600Hz) and temporal extents (10^-2^-10^2^ s)^1^. Identifying the frequency, temporal location, duration, and magnitude of finite oscillation packets with high precision is a significant challenge.

Time-frequency analysis of digitized signals is traditionally performed using the short-time Fourier transform (STFT) ^2^, which computes Fourier spectra on successive sliding windows. Long windows provide good frequency resolution but poor temporal resolution, while short windows increase temporal resolution at the expense of frequency resolution. This is known as the Heisenberg-Gabor uncertainty principle ^3^ or the Gabor limit ^4^, i.e. one cannot simultaneously localize precisely a signal in both time and frequency. Importantly, this limit applies to a single measurement. Frequency resolution is proportional to window size, as defined by the Rayleigh frequency ^5,6^. Therefore, shortening the window to gain temporal resolution leads to a degradation of frequency resolution (Fig. 1A, left).

**Fig. 1.**
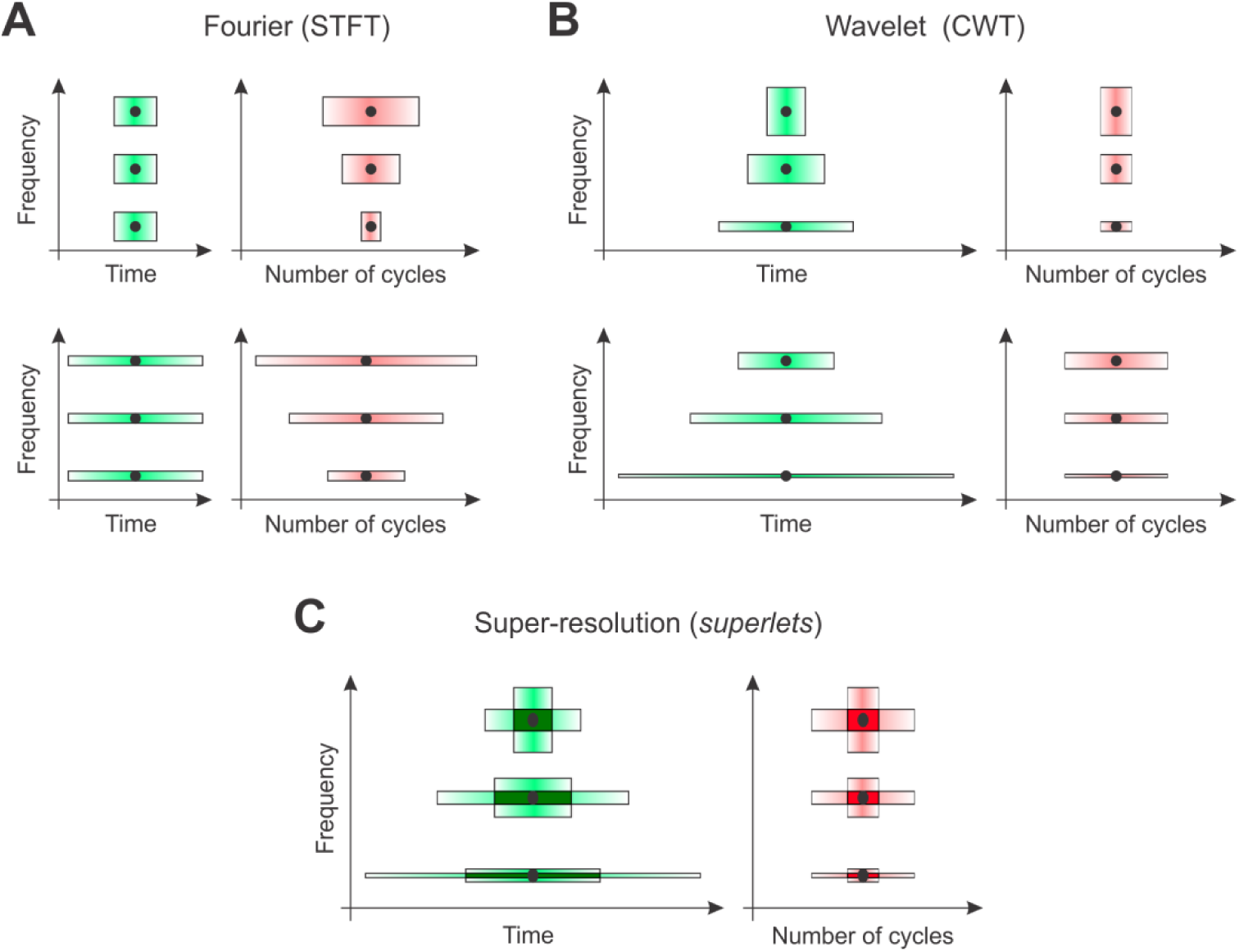
Sketch of time-frequency uncertainty in Fourier (STFT), wavelet (CWT), and *superlet* (SLT) analysis. (**A**) Time and frequency resolution of the STFT for a short (top) and wide (bottom) window at three different frequencies. Temporal resolution is expressed in time (left) or in oscillation cycles at the target frequency (right). (**B**) Same as in (**A**) but for wavelets (CWT). Here, the number of cycles is fixed across the spectrum but the spanned temporal window decreases with frequency increase. (**C**) *Superlets* of order 2 (SLT). Time-frequency super-resolution is achieved by combining short, low-bandwidth wavelets, with longer, high-bandwidth wavelets.

For a given window size, the STFT has fixed frequency resolution but its temporal precision relative to period decreases with increasing frequency (Fig. 1A, right). To overcome this limitation multi-resolution techniques have been introduced, like the continuous-wavelet transform (CWT). The CWT provides good temporal localization by compression/dilation of a mother wavelet as a function of frequency ^7^. The most popular wavelet for time-frequency analysis is the Morlet ^8,9^, defined as a plane wave multiplied by a Gaussian envelope (see Fig. S1):

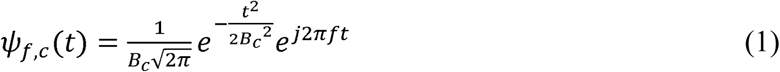

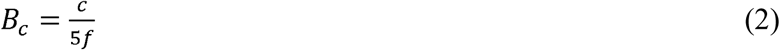

where, *f* is the central frequency, *c* is the number of cycles of the wavelet, *B_c_* is the bandwidth (in Hz^-1^ = s) or variance of the wavelet ^10^. A Morlet wavelet with higher bandwidth contains more cycles, is wider in time, but has a narrower frequency response. Here we will use Morlet wavelets for time-frequency analysis, but other choices are also possible.

The CWT localizes well the oscillation packets in time, but trades in frequency resolution as frequency increases ^11,12^ (Fig. 1B). Neighboring high frequencies cannot be distinguished, i.e. the representation is redundant across wavelets with close central frequencies in the high-range. For this reason, analyses are often performed using a diadic representation, as in the discrete wavelet transform (DWT), where frequencies are represented as powers of 2 ^12,13^. This representation however resolves very poorly the high-frequencies.

Both the STFT and CWT (or DWT) have significant limitations. The STFT provides good frequency resolution but poor temporal resolution at high frequencies, while the CWT maintains a good temporal resolution throughout the spectrum but degrades in frequency resolution and becomes redundant with increasing frequency. This time-frequency uncertainty hampers analysis of neuronal signals, which have rich time-frequency content ^14,15^.

To overcome the limitations of the STFT it has been proposed to combine Fourier-based spectrograms obtained with a short and a long window ^16^, or with a set windows with varying sizes ^17^. This technique was termed super-resolution ^18,19^ because it can localize oscillation packets simultaneously in both time and frequency better than it is possible with any single spectrogram. A similar idea is applied in super-resolution methods used in imaging ^20,21^.

To increase resolution, the information from multiple spectrograms can be combined by computing their geometric mean ^16^, which is equivalent to the minimum mean cross-entropy (MMCE) ^17,22^, which is optimal with respect to an entropic criterion ^17^. Here, we use the minimum cross-entropy technique in combination with wavelets to introduce a novel approach that reveals a sharper localization of oscillation packets than can be achieved with STFT, CWT, or the Fourier-based MMCE.

### Superlets

In structured illumination microscopy (SIM) one uses a set of known illumination patterns ^21^ to obtain multiple measurements that are combined to achieve super-resolution. The time-frequency super-resolution technique proposed here employs multiple wavelets to detect localized time-frequency packets.

The method can be formalized as follows. A base wavelet, e.g. Morlet with a fixed number of cycles, provides multi-resolution in the standard sense, with constant relative temporal resolution but degrading frequency resolution (increased redundancy) as the central frequency of the wavelet increases. By increasing the bandwidth of the wavelet (more cycles), one increases frequency resolution (Fig. 1B) but loses temporal resolution. To achieve super-resolution we propose to combine short wavelets having high temporal resolution (small number of cycles, low bandwidth) with longer wavelets, having high frequency resolution (larger number of cycles, lower temporal resolution) (Fig. 1C).

A “*superlet*” (SL) is defined as a set of Morlet wavelets with a fixed central frequency, *f*, and spanning a range of different cycles (bandwidths):

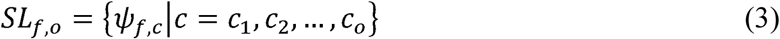

where, *o* is the “order” of the *superlet*, and *c_1_*, *c_2_*,…, *c_o_* are the number of cycles for each wavelet in the set. A *superlet* of order 1 is a single (base) wavelet with *c_1_* cycles. In other words, a *superlet* is a finite set of *o* wavelets spanning multiple bandwidths at the same central frequency, *f*. The order of the *superlet* represents the number of wavelets in the set. The number of cycles defining the wavelets in the *superlet* can be chosen *multiplicatively* or *additively*. In a multiplicative *superlet*, *c_i_* = *i · c_1_*, whereas in an additive *superlet c_i_* = *c_1_ + i* – 1, for *i* = 2,…, *o*.

We define the response of a *superlet* to a signal, *x*, as the geometric mean of the responses of individual wavelets in the set:

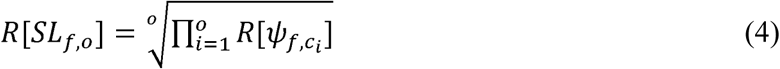

where, *R*[*ψ_f,ci_*] is the response of wavelet *i* to the signal, i.e., the magnitude of the complex convolution (for complex wavelets, such as Morlet):

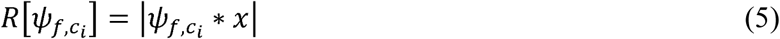

where, * is the convolution operator and *x* the signal. The *superlet* is an estimator of the magnitude of oscillation packets present in the signal at the central frequency, *f*, of the *superlet*. To estimate power, the response of the *superlet* is simply squared. We will show that, while increasing frequency resolution locally, the *superlet* does not significantly lose time resolution.

The *superlet* transform (SLT) of a signal is computed analogously to the CWT, except that one uses *superlets* instead of wavelets. A SLT with *superlets* of order 1 is the CWT. As will be shown next, the SLT with orders > 1 is a less redundant representation of the signal than the corresponding CWT.

### Adaptive superlets

At low central frequencies, single wavelets (i.e., *superlets* of order 1) may provide sufficient time-frequency resolution. Indeed, the CWT is less redundant at low than at high frequencies ^12^. Adaptive *superlets* (ASL) adjust their order to the central frequency to compensate the decreasing bandwidth with increasing frequency. In an adaptive *superlet* transform (ASLT), one starts with a low order for estimating low frequencies and increases the order as a function of frequency to achieve an enhanced representation in both time and frequency across the entire frequency domain, as follows:

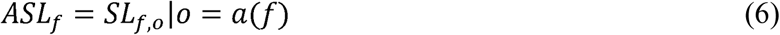

where, *a(f)* is a monotonically increasing function of the central frequency, having integer values. A simple choice is to vary the order linearly:

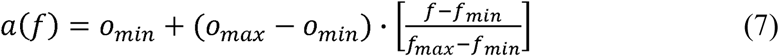

where, *o_min_* is the order corresponding to the smallest central frequency, *f_min_*, *o_max_*is the order corresponding to the largest central frequency, *f_max_*, in the time-frequency representation, and [] is the nearest integer (round) operator. We recommend using the ASLT when a wide frequency range needs to be resolved, and the SLT for narrower bands.

An implementation of *superlets* and test code can be downloaded here: http://muresanlab.tins.ro/sources/superlets/superlets.zip.

## Results

We will first illustrate the basic principle behind *superlets* by considering a known set of packets composed of 7 sinusoidal cycles. A target oscillation packet, *T*, is composed of a finite number of cycles at a target central frequency. We define two additional oscillation packets: a temporal neighbor *NT*, having the same frequency, but shifted in time with a temporal offset Δ*t*, and a frequency neighbor *NF*, at the same location in time, but shifted with a frequency offset Δ*f* (Fig. 2A, top). For convenience, all three packets have a magnitude of 1.

**Fig. 2.**
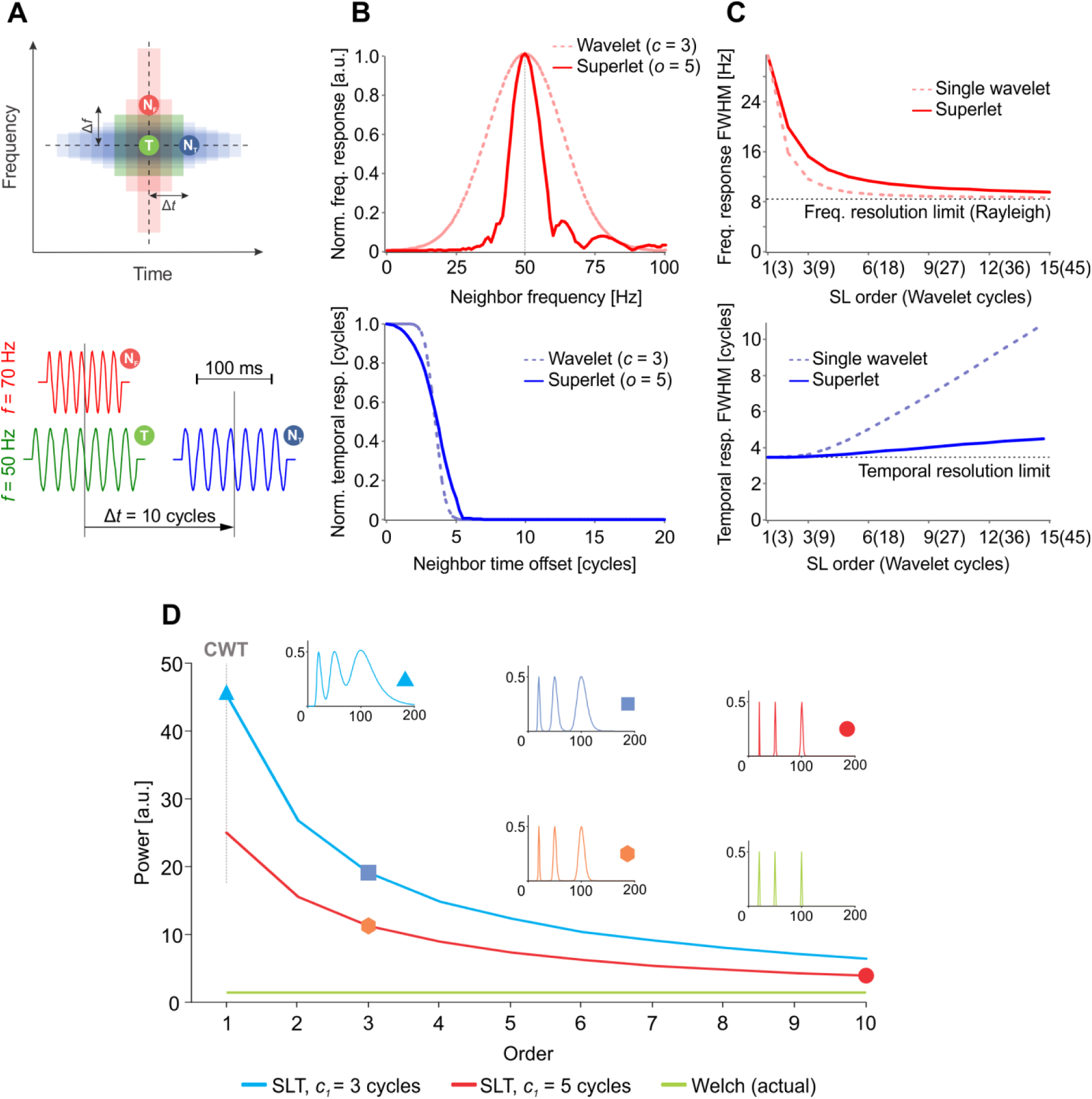
The principle behind *superlets* and redundancy of spectral representation. (**A**) Test setup where a target oscillation packet, *T*, is contaminated by frequency and time neighbors (*N_F_* and *N_T_*). Top: using a wavelet with small number of cycles enables good time separation but has poor frequency resolution (red), while wavelets with many cycles enable good frequency separation but suffer from temporal contamination (blue). Bottom: a particular instantiation with packets of 7 cycles having: target frequency 50 Hz, neighbor frequency 70 Hz, neighbor time offset 10 cycles. (**B**) Target contamination in frequency by *N_F_* (top) and in time by *N_T_*(bottom). Contamination is measured as the normalized response (magnitude) of a single wavelet (*c* = 3) or a multiplicative *superlet* (*c_1_*= 3; *o* = 5) at the time-frequency location of the target (without the target being present) to *N_F_* with various frequencies (top) or *N_T_* with various time offsets (bottom). (**C**) Frequency (top) and time (bottom) *superlet* resolution measured as the half-width of the frequency and time peak in (**B**), respectively, as a function of the order of a multiplicative *superlet* (line). The same is shown for the longest wavelet in the *superlet* set (dotted line). The frequency resolution limit is the Rayleigh frequency of *T* with Gaussian windowing. The temporal resolution limit is half the size of *T* (3.5 cycles). (**D**) A long signal composed of 3 summed unitary amplitude sine waves has an average power of 1.5 (green). Two *superlet* transforms (SLT) using multiplicative *superlets* with *c_1_* = 3 (blue) and 5 (red) give an increasingly sharper representation of the higher frequencies, as their order is increased. Sharper representations signify less redundancy.

An example instantiation of this scenario is shown in Fig. 2A, bottom, for a target frequency of 50 Hz in a signal sampled at 1 kHz. We next evaluated how the presence of *NF* or that of *NT* influences the estimation at the location of *T*. In other words, without *T* being present, we systematically moved *NF* in frequency or *NT* in time and computed their contribution (leakage) to the estimate at the time-frequency location of *T* (Fig. 2B). As estimators, we initially considered a wavelet with *c*=3 cycles and a multiplicative *superlet* with *c_1_*=3 and *o*=5. The bandwidth of the wavelet was poor, with a broad frequency response around the target frequency of *T*, indicating that *T* was hard to distinguish from *NF* over a large frequency domain (Fig. 2B, top). By contrast, the *superlet* significantly sharpened the frequency response, reducing frequency cross-talk between *NF* and *T.* Along the temporal dimension, when *NT* was shifted in time away from the target’s location (time offset 0), the response of both the wavelet and the *superlet* dropped sharply after half the size of the target packet (3.5 cycles) (Fig. 2B, bottom). This indicates that, while significantly increasing frequency resolution, the *superlet* did not induce a significant loss of temporal resolution.

To investigate how the *superlet* achieves high frequency resolution without losing temporal resolution we quantified these two properties by evaluating the full width at half maximum (FWHM) of the frequency and temporal responses measured at *T* and induced by *NF* and *NT*, respectively (Fig. 2C). We varied the order of the *superlet* and compared its response to the one of the longest wavelet in its corresponding set (*c_o_*, see eq. 3). As the order was increased, both the largest wavelet (with highest bandwidth) and the *superlet* approached the frequency resolution limit (Rayleigh frequency corresponding to the Gaussian-windowed oscillation packet) (Fig. 2C, top). By contrast, while the single wavelet’s temporal resolution decreased rapidly by increasing its number of cycles, the temporal resolution of the *superlet* degraded considerably slower (Fig. 2C, bottom). These results indicate that, as its order is increased, a *superlet* nears the theoretical frequency resolution possible for a limited duration oscillation packet (Rayleigh frequency) while maintaining a significantly better time resolution than a single, long wavelet. By approaching the theoretical frequency resolution limit without losing temporal resolution the *superlet* achieves time-frequency super-resolution.

The CWT provides a representation of the signal that is increasingly redundant for higher frequencies ^11,12^ because the frequency response of wavelets becomes broader as the number of samples per oscillation cycle decreases. The SLT (and ASLT) decreases the redundancy of the representation with increasing order of the *superlets*. Figure 2D depicts the average power measured over a long signal composed of three frequency components (20, 50, and 100 Hz) with unitary amplitude. The average power in a perfect energy-conserving transform should be 1.5 (Fig. 2D, green). We used two types of *superlets* with base cycles *c_1_*= 3 and 5, and progressively increased their order while computing the SLT and collapsing it in time (Welch-like). Order 1 corresponded to the CWT and, as the order was increased, the redundancy in the representation of high frequencies was reduced (Fig. 2D, insets) and the average power across the spectrum approached that of an energy-conserving transform (e.g., Fourier). Importantly, *superlets* with larger base cycles provide a less redundant representation than those with smaller number of base cycles (compare Fig. 2D red with Fig. 2D blue), albeit at the expense of decreased temporal resolution (see below).

In another test, we generated a signal as a sum of multiple time-frequency packets (Fig. 3A), as follows. Three target packets of 11 cycles were generated at target frequencies of 20, 40, and 60 Hz. For each target, a neighbor in frequency (+10 Hz) and a neighbor in time (+12 cycles) were added to the signal. Due to constructive-destructive summation, a clear modulation of magnitude is visible where the target was summed with its frequency neighbor – this is equivalent to amplitude modulation (AM) whereby two sideband frequencies sum up to give rise to an amplitude modulated central frequency. Locally, the correct time-frequency representation of this phenomenon should reveal corresponding bursts of magnitude (or power) at the two summed frequencies (i.e., locally the signal looks like it is modulated in amplitude and it is composed of two AM sideband component; thus, locally, both interpretations are correct at the same time). We computed the time-frequency power representation of the signal using Blackman-windowed Fourier (STFT), wavelets (CWT), and adaptive additive superlets (*o* = 1:30; order varied linearly from 1@10 Hz to 30@75 Hz) (see Fig. 3).

**Fig. 3.**
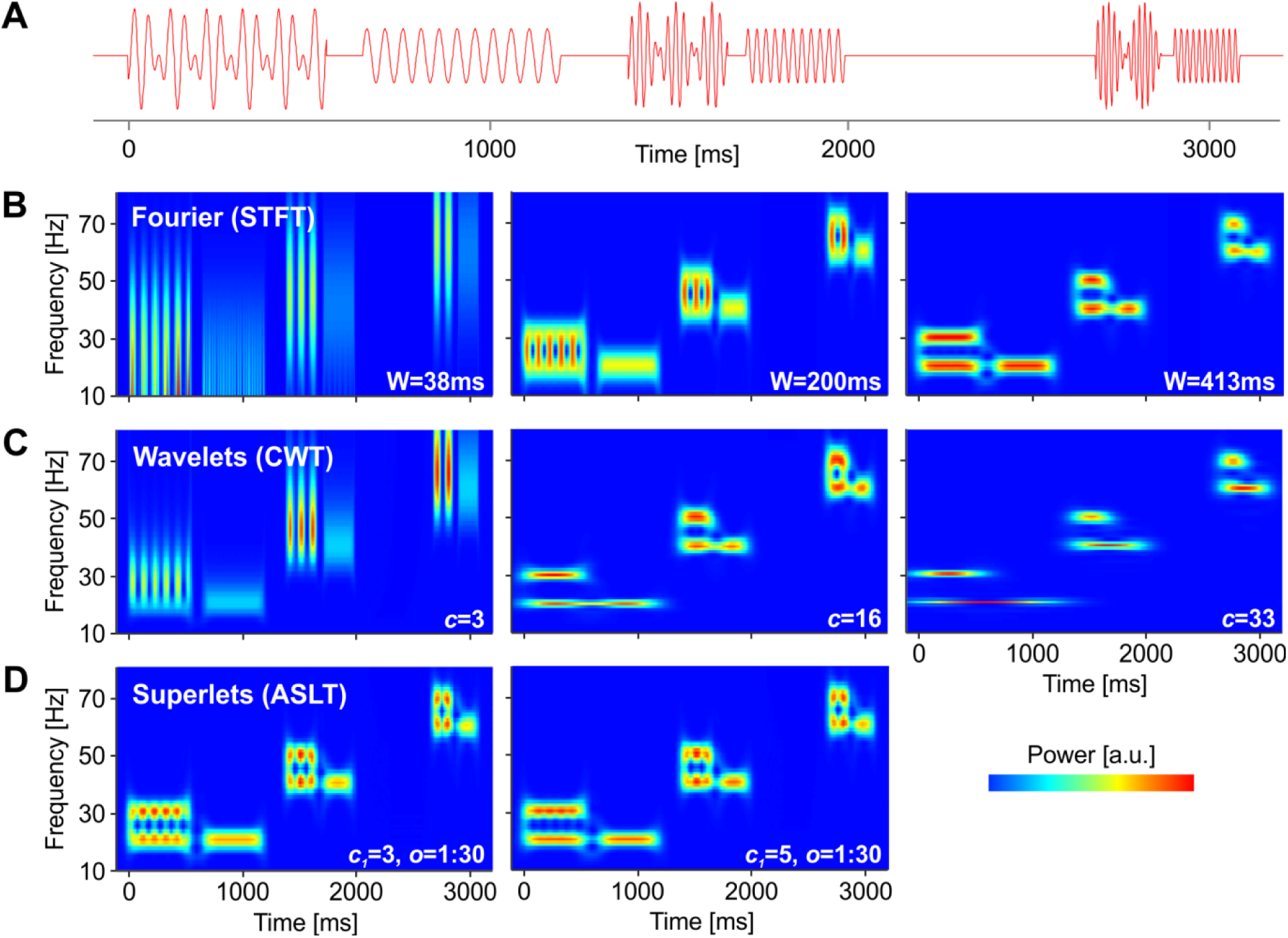
Evaluation of time-frequency resolution on a known signal structure. (**A**) Test signal containing 3 target packets with 11 cycles at 20, 40, and 60 Hz, each accompanied by a neighbor in frequency (+10 Hz) and a neighbor in time (+12 cycles). The signal sampling rate is 1024 samples/s. (**B**) Time-frequency power representation of the signal using STFT (Blackman window) and window size 38, 200, and 413ms (roughly matching the size of a single wavelet with 3, 16, 33 cycles at the largest target frequency). (**C**) Same as in (**B**) but using Morlet wavelets with increasing number of cycles. (**D**), Same as (**B**) and (**C**) but using adaptive additive *superlets* with linearly varying order from *o* = 1, for 10 Hz, to *o* = 30, for 75 Hz, and with different number of base cycles (*c_1_* = 3, left; *c_1_*= 5, right).

At the location of frequency neighbors, the STFT with various window sizes revealed either the temporal modulation (Fig. 2B, left) or the two frequencies (Fig 2B, right), but it was unable to fully segregate time and frequency, in spite of an “optimized” intermediate window size (Fig. 2B, center). A similar conclusion was reached with a CWT using increasing number of wavelet cycles (increasing bandwidth; Fig. 2C), with the difference that the CWT provided better frequency resolution in the low frequency range. By contrast, adaptive *superlets* (ASLT) provided a faithful local representation with high resolution in both time and frequency across the entire spectrum (Fig. 3D). Increasing the number of base cycles (*c_1_*) had the effect of further increasing frequency precision, albeit at the cost of losing some temporal resolution at the low frequencies.

We next used *superlets* to analyze brain signals (EEG) recorded from humans in response to visual stimuli representing objects (deformable dot lattices) ^23^ (see Materials and Methods). Because EEG signals are strongly affected by the filtering properties of the skull and scalp, having a pronounced 1/*f* characteristic ^24,25^ that masks power in the high frequency range, we have baselined spectra to the pre-stimulus period ^26^. The time-frequency power spectrum of the occipital signal over the Oz electrode was estimated using STFT, CWT, and ASLT (Fig. 4A). The STFT window was chosen to optimize the representation in the gamma range (> 30 Hz), while the number of cycles for the CWT was chosen to maximize temporal resolution. The STFT provided a poor resolution in the low frequency range (Fig. 4A, top), while the CWT showed good temporal resolution but poor frequency resolution for higher frequencies (Fig. 4A, middle). On the same data, the ASLT provided sharp time-frequency resolution across the whole frequency range and revealed fine details that could not be resolved by the other methods (Fig. 4A, bottom).

**Fig. 4.**
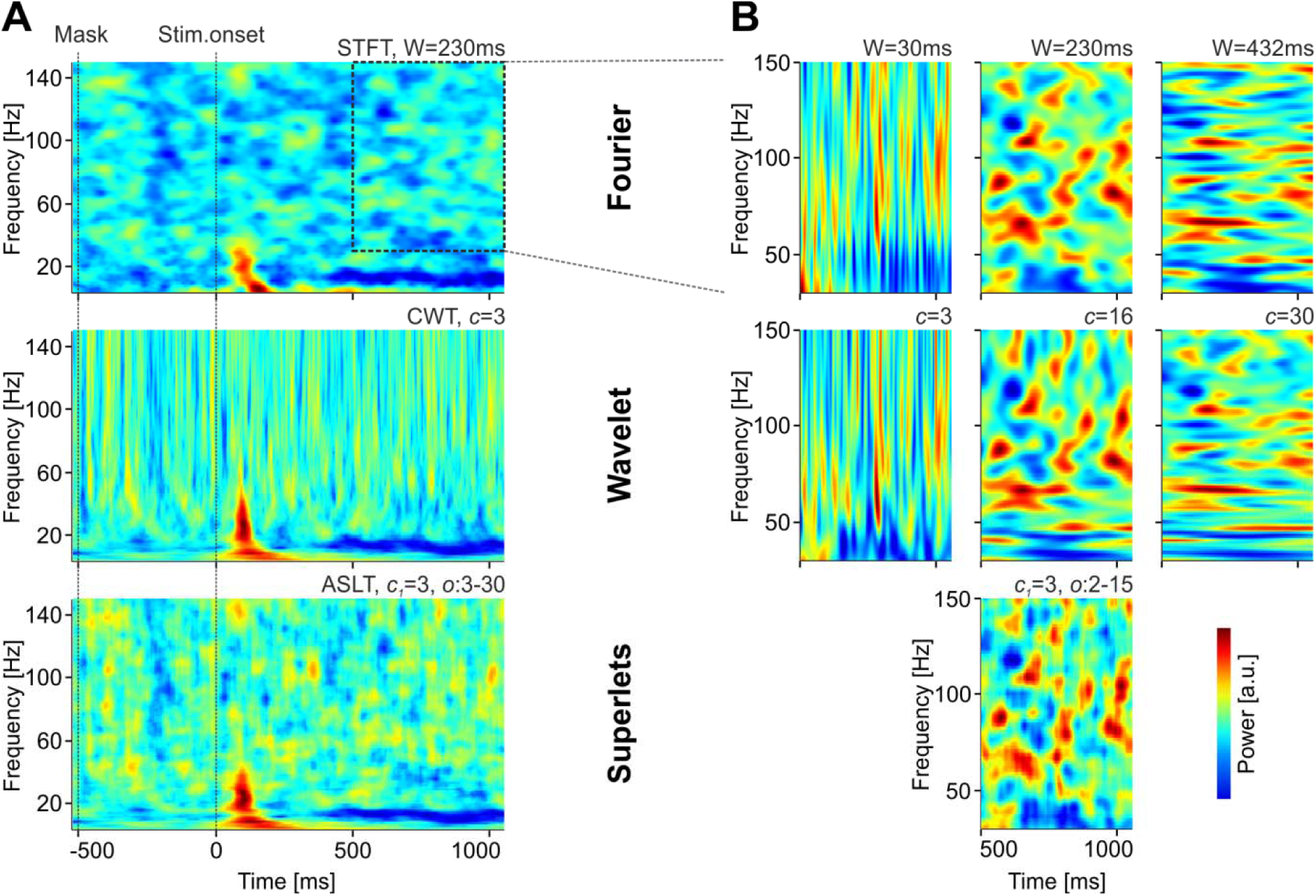
Time-frequency analysis of the EEG signal recorded over occipital electrode Oz. (**A**) Global time-frequency power spectrum around stimulus onset computed using Fourier analysis (STFT; top), wavelets (CWT; middle), and adaptive additive *superlets* (ASLT; bottom). Power scale is logarithmic. (**B**) Zoom-in analysis over the gamma frequency band (30-150Hz) using STFT with various windows (top), CWT with different number of Morlet cycles (middle), and adaptive multiplicative *superlets* (bottom). Power scale is linear. All analyses are baselined to 1.5s pre-stimulus period.

We next zoomed in on the gamma frequency range, which poses particular challenges for time-frequency analysis ^27–30^. The Fourier window (Fig. 4B, top row) and the number of wavelet cycles (Fig. 4B, middle row) were varied to optimize the temporal (left) or frequency (right) precision, or a trade-off between the two (middle). *Superlets* (Fig. 4B, bottom) shared the major features with the other representations but provided time-frequency details that could not be simultaneously resolved by any of the latter.

*In vivo* electrophysiology signals are recorded at much higher sampling rates than EEG (32 kHz compared to 1 kHz), offering the opportunity to observe time-frequency components with higher resolution in local-field potentials (LFP) than in EEG. We next focused on LFPs recorded from mouse visual cortex during presentation of drifting sinusoidal gratings (see Materials and Methods). LFPs suffer from the 1/*f* issue significantly less than EEG and therefore baselining is typically not necessary for their analysis. We computed the time-frequency representation of an LFP signal using the STFT (Fig. 5A, top), CWT (Fig. 5A, bottom), and ASLT (Fig. 5A, middle) around the presentation of the visual stimulus (drifting grating at 45°) and averaged it across 10 presentations (trials). As was the case for EEG data, adaptive *superlets* provided the best time-frequency representation across the entire analyzed spectrum. They revealed 45 Hz gamma bursts induced by the passage of the grating through the receptive fields of cortical neurons ^31^ and resolved many details in both the low and high frequency range.

**Fig. 5.**
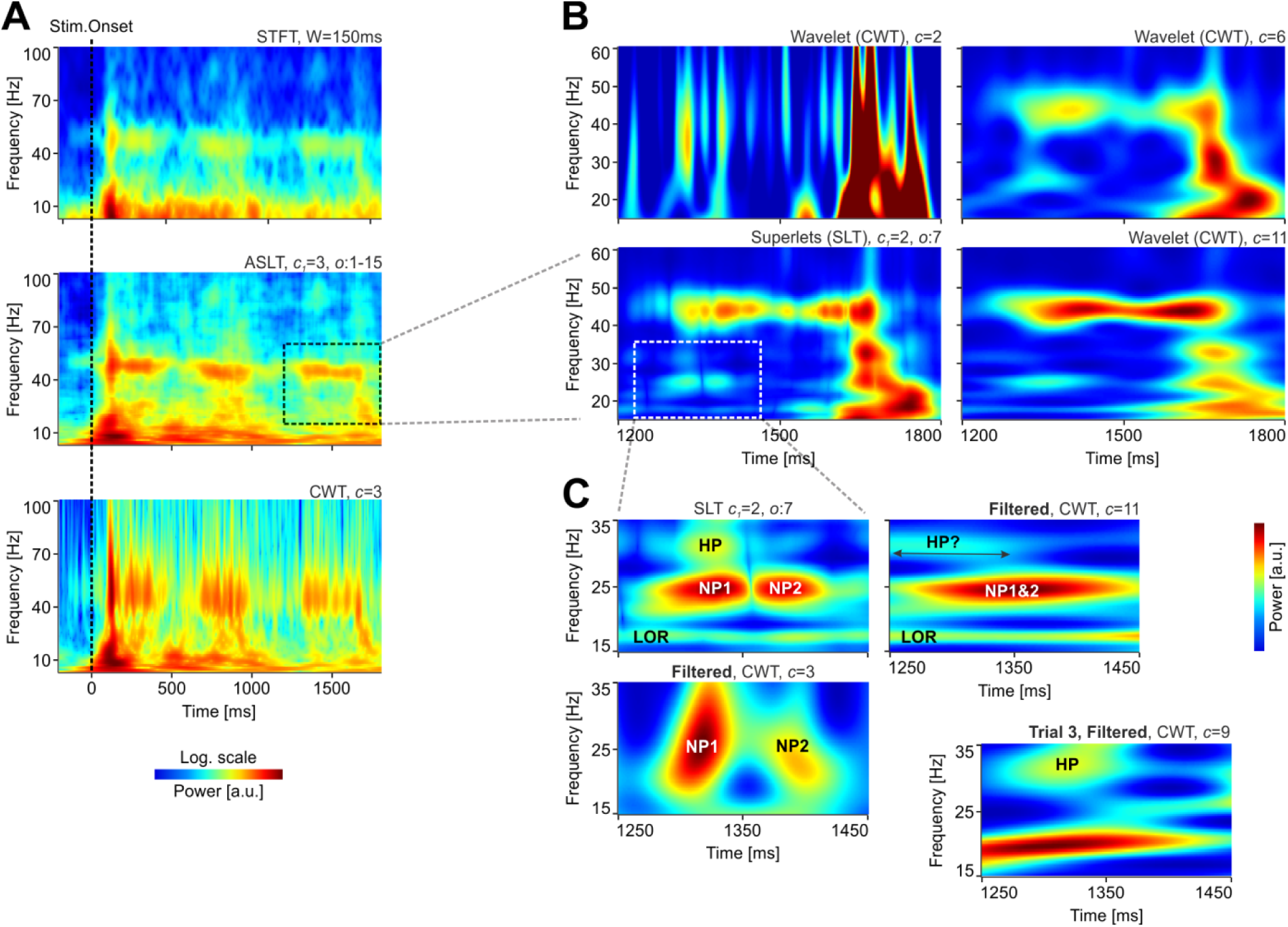
Time-frequency analysis of acute electrophysiology data recorded in mouse visual cortex. (**A**) Fourier (STFT; top), adaptive multiplicative *superlets* (ASLT; middle), and wavelet power spectra (CWT; bottom) around stimulus onset. (**B**) Zoom-in on a gamma burst induced by the passage of the grating through the receptive field of cortical neurons. The SLT used multiplicative *superlets* of order 7 and *c_1_* = 2, optimized to provide high temporal and frequency resolution (bottom-left). By comparison, individual walevets optimized for time (top-left), frequency (bottom-right) or a compromise between the two (top-right) cannot reveal all the details evidenced by the *superlet*. (**C**) Further zoom-in on a detail provided by the *superlet* (top-left). Tuned wavelets on 10-40Hz band-passed data indicate roughly the presence of the temporal (left-bottom) and frequency (top-right) components. The location of a high-frequency packet (HP) cannot be determined by wavelet analysis in the average time-frequency spectrum, but is recovered by single-trial analysis, indicating that *superlets* can correctly reveal very fine time-frequency details, which are smeared out in the average spectra by other methods. No baselining used. The power scale is logarithmic in (**A**), but linear in (**B**) and (**C**).

The true power of *superlets* was, however, revealed when we zoomed in on a compact gamma burst induced by the passage of the grating (see Fig. 5B). *Superlets* were computed with a base cycle *c_1_* = 2, to maximize temporal resolution, and we used a fixed multiplicative order of 7 (SLT). The SLT provided very fine temporal and frequency details, whose presence in the signal was validated by computing the local CWT optimized for time (*c* = 2), frequency (*c* = 11), or a tradeoff between time and frequency (*c* = 6). The components seen in the *superlet* representation could be inferred from these multiple wavelet representations but none of the latter was able to simultaneously reveal all the time-frequency details (Fig. 5B).

To push the envelope, we further explored a time-frequency detail revealed by *superlets* (Fig. 5B, left-bottom and Fig. 5C), composed of a lower ongoing rhythm at ∼17.5 Hz (LOR), two time neighboring packets at 24.5 Hz (NP1 and NP2), and one higher frequency packet at ∼31 Hz (HP) (Fig. 5C, top-left). To determine if these features were actually present in the signal over the 10 trials, we narrow-band filtered the signal (bidirectional IIR, order 3, band-pass 10-40 Hz) to remove frequency contamination plaguing the wavelet estimates in the gamma range. This enabled us to validate the presence of the time-frequency packets using narrow (*c* = 3) and wider (*c* = 11) wavelets (Fig. 5C, bottom-left and top-right). However, while the frequency of HP could be identified, its clear temporal location could not be established with the CWT, irrespective of the parameters of the wavelet (Fig. 5C, top-right). We suspected that this may originate from averaging over 10 trials such that time/frequency smearing of the long/short CWT could hide this detail. Indeed, we found that HP was expressed clearly in at least one of the trials in the set (Fig. 5C, bottom-right). Thus, all the packets in the time-frequency detail revealed by *superlets* were actual features in the signal and, in addition, the time-frequency concentration provided by *superlets* was able to reveal bursts expressed at single-trial level. The latter, could not be identified by other methods because they were averaged out due to the time/frequency smearing in the wavelet or Fourier representations (see also Fig. S2).

One of the hardest problems in the time-frequency analysis of brain signals is to resolve weaker neighboring rhythms when strong, dominating oscillation bursts are present in a certain frequency band. For example, oscillations may occur simultaneously in the beta and gamma-bands and these are often hard to distinguish ^28^. Figure 6 shows an example analysis on the same data from Fig. 5 but in response to a different grating direction (135°) and focused on the post stimulus-onset part of the response. All methods were able to reveal the strong gamma bursts occurring in response to the drifting grating (Fig. 6A). The STFT displayed strong leaked-in power from the lower frequencies (Fig. 6A, top) while the CWT had markedly poor frequency resolution (Fig. 6A, middle). As before, the SLT provided the best time-frequency resolution (Fig. 6A, bottom). Importantly, the latter also suggested the presence of time-frequency structure in the beta band (12-30 Hz).

**Fig. 6.**
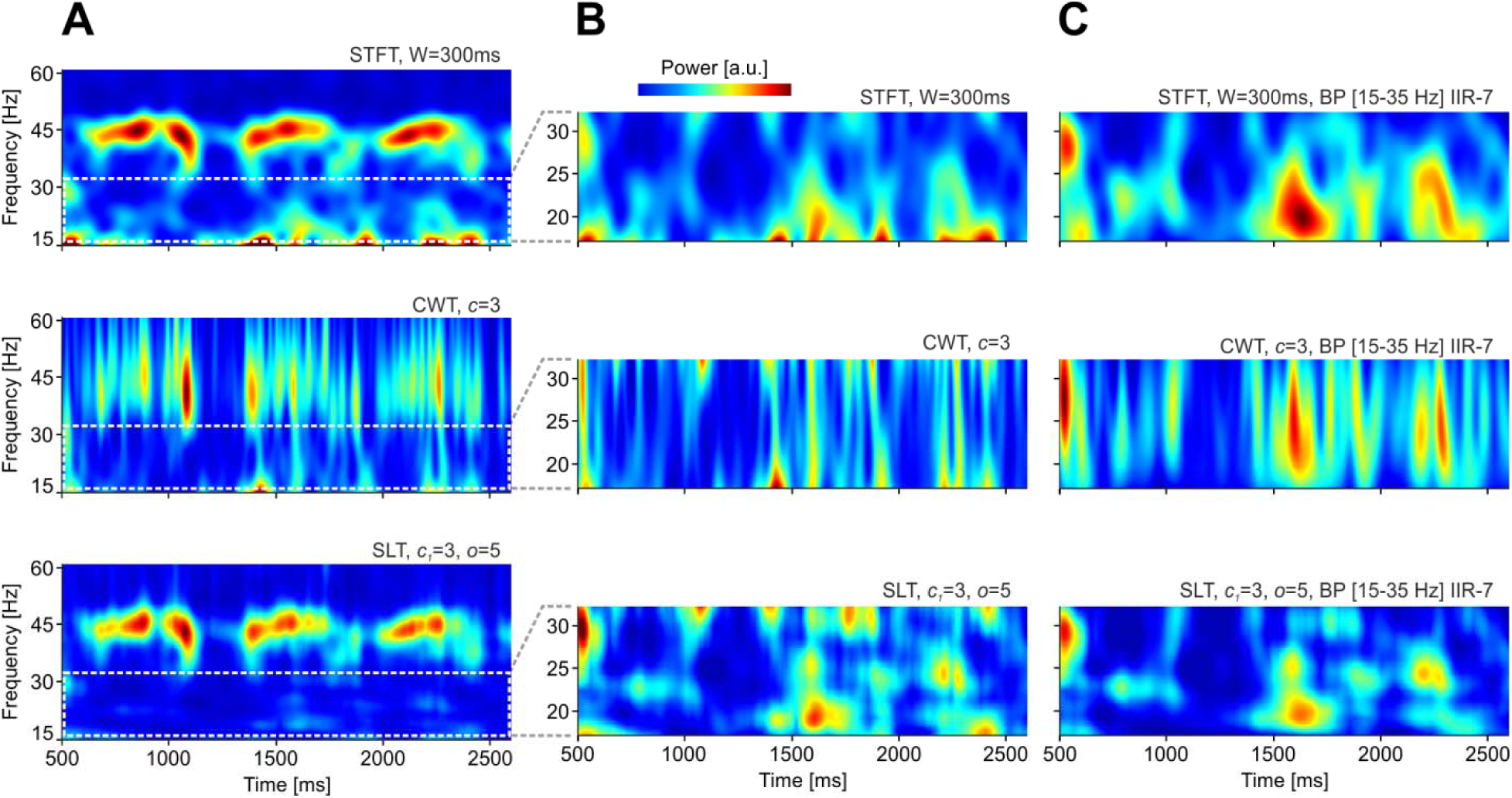
Effect of averaging time-frequency spectra. (**A**) Fourier (STFT; top), wavelet power spectra (CWT; middle), multiplicative *superlets* (SLT; bottom) computed for the beta- and low-gamma bands (15-60Hz) and averaged across 10 trials. Stimulus onset was excluded from analysis to focus on the fast oscillations induced by the passage of the drifting grating. (**B**) Zoom-in on the beta-band. (**C**) Same as in (**B**) but computed after the signal was band-pass filtered in 15-35Hz using a bidirectional Butterworth IIR filter of order 7. No baselining used. The power scale is linear.

When we zoomed in on the beta band, the SLT revealed a rich time-frequency structure with multiple packets located at various frequencies and temporal offsets (Fig. 6B, bottom). By contrast, the STFT (Fig. 6B, top) and CWT (Fig. 6B, middle) were unable to properly resolve this frequency band, suffering from strong power leakage from below and above the band. When the signal was narrow-band filtered in the 15-35Hz range, the representation in the STFT (Fig. 6C, top) and the CWT (Fig. 6C, middle) changed dramatically, and became more similar to the SLT, roughly revealing the same time-frequency structure. By contrast, the SLT maintained a very similar representation after filtering, implying that it was the only method correctly capturing the time-frequency structure of the beta band. Thus, of all the tested time-frequency estimation techniques, *superlets* provided the most accurate representation of the beta band, with minimal interference from bursts in the frequency bands below and above it.

The closest time-frequency super-resolution method to *superlets* is the Fourier-based MMCE ^22^. The latter employs a set of windows with increasing size and combines the resulting spectrograms geometrically. We next compared how *superlets* faired with respect to the established MMCE method. Fig. 7 depicts a comparison of the two methods when applied to EEG data (same experiment like in Fig. 4, different subject). Signals were aligned to either to the presentation of the visual stimulus (onset) or the response of the subject (button press) and analyses were grouped across 3 conditions, as a function of the response of the subject (object seen, uncertain, nothing seen) ^23^. The MMCE, computed with a set of 7 windows, spanning from 30 ms to 700 ms, performed rather poorly, especially on the stimulus onset aligned data (Fig. 7A). While frequency resolution remained constant, relative time resolution degraded with increasing frequency. Also, a marked effect was the masking / “dilution” of higher frequency packets when powerful low-frequency components are present. *Superlets* did not suffer from these problems and revealed a rich time-frequency structure, independently of how the data was aligned (Fig. 7B). All analyses revealed interesting associations between alpha, beta, and gamma activity and the experimental condition, but *superlets* provided a sharper picture. The increase in gamma bursts expression and in their frequency was associated with the perceptual process, while increased ongoing beta was more sustained in association to response certainty in this particular subject. Alpha was in general suppressed during stimulus processing.

**Fig. 7.**
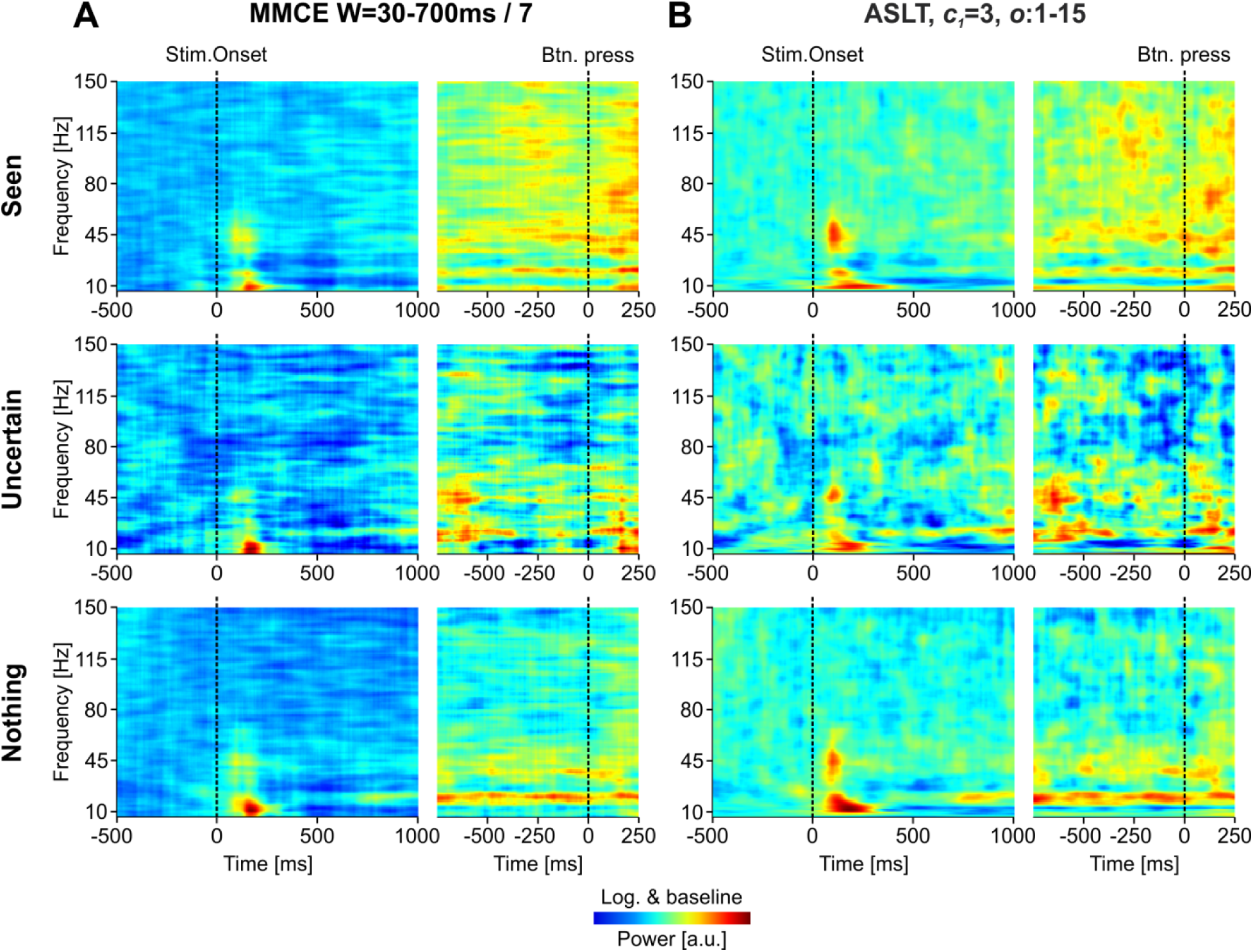
Comparison between minimum mean cross-entropy (MMCE) and *superlets* on time-frequency analysis across 15 occipital EEG electrodes. (**A**) MMCE using 7 windows that span 30-700 ms. (**B**) Adaptive multiplicative *superlets* with orders spanning 1-15 and *c*_1_ = 3 cycles. For each analysis, signals were aligned to stimulus onset (left) or subject response (right). Data were log-ed and baselined to pre-stimulus period ^26^. Color scales were maintained identical across conditions (along columns) to facilitate comparisons.

We next compared the two methods on electrophysiology data. Fig. 8 shows results on the same data as in Fig. 6, but for a larger time and frequency range. As was the case for EEG data, the MMCE displayed a progressive “dilution” of oscillation packets with increasing frequency (Fig. 8A). In addition, the MMCE provided poorer frequency resolution for the low frequencies, i.e. frequency resolution did not adapt across the spectrum. By contrast, adaptive *superlets* did not display the “dilution” effect and provided robust details across the entire frequency range (Fig. 8B). To investigate this “dilution” effect, we optimized both the MMCE and *superlets* to estimate a limited gamma burst induced by the drifting grating (Fig. 8A-C, inset 1). In this limited range, both methods provided rather identical results, thus validating the fact that *superlets* provide locally consistent results with established methods. However, when we expanded the analysis window to incorporate a progressively larger frequency range (Fig. 8A-C, insets 2 and 3), the MMCE displayed a marked dilution of the higher frequencies while the *superlets* did not.

**Fig. 8.**
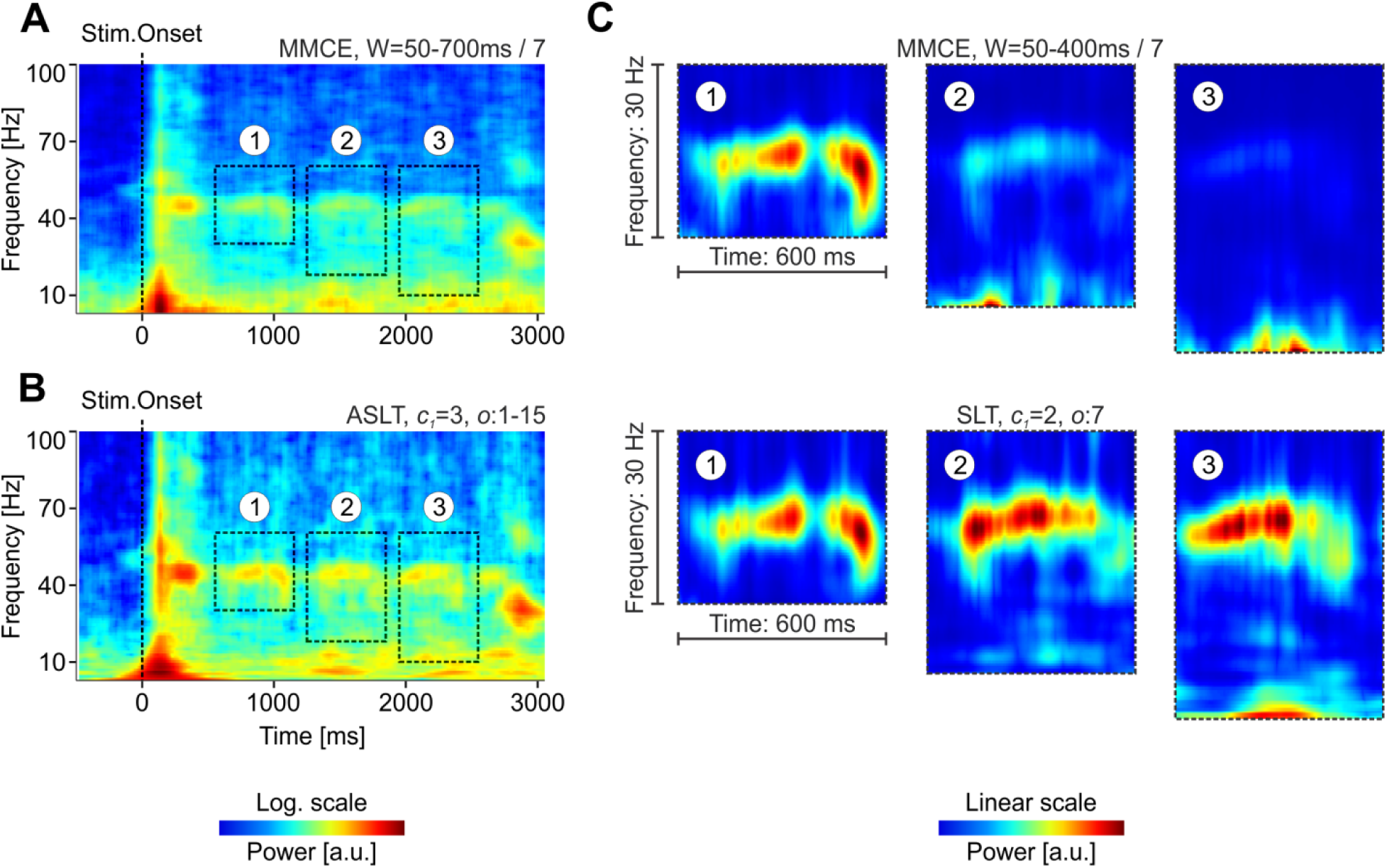
Comparison between minimum mean cross-entropy (MMCE) and *superlets* on time-frequency analysis of *in vivo* electrophysiology signals. (**A**) MMCE using 7 windows that span 50-700 ms. (**B**) Adaptive multiplicative *superlets* with orders spanning 1-15 and *c*_1_ = 3 cycles. For each analysis, signals were aligned to stimulus onset (left) or subject response (right). (**C**) Analyses on restricted windows with various locations and sizes [see insets in (**A**) and (**B**)] using MMCE (top) and *superlets* (bottom). The power scale is logarithmic in (**A**) and (**B**), to facilitate exploration of a large spectral range, while it is linear in (**C**), to enable precise quantitative comparison.

During cognitive processes it is expected that gamma bursts are scattered in time and frequency in a single-trial dependent fashion. Therefore, we evaluated the ability of *superlets* to discover gamma bursts expressed in single trials. Three packets of 40, 80, and 120Hz were inserted into only one of the 84 trials (Fig. 9A) recorded over the Pz electrode in condition “Nothing” of the dataset shown in Fig. 7. We computed the averaged time-frequency spectrum over the 84 trials using STFT (Fig. 9B), CWT (Fig. 9C), MMCE (Fig. 9D), and multiplicative SLT (Fig. 9E). Only *superlets* were able to robustly detect the presence of all three packets. The latter also revealed a rich time-frequency structure in the upper frequencies (>50Hz), which the other methods mostly missed.

**Fig. 9.**
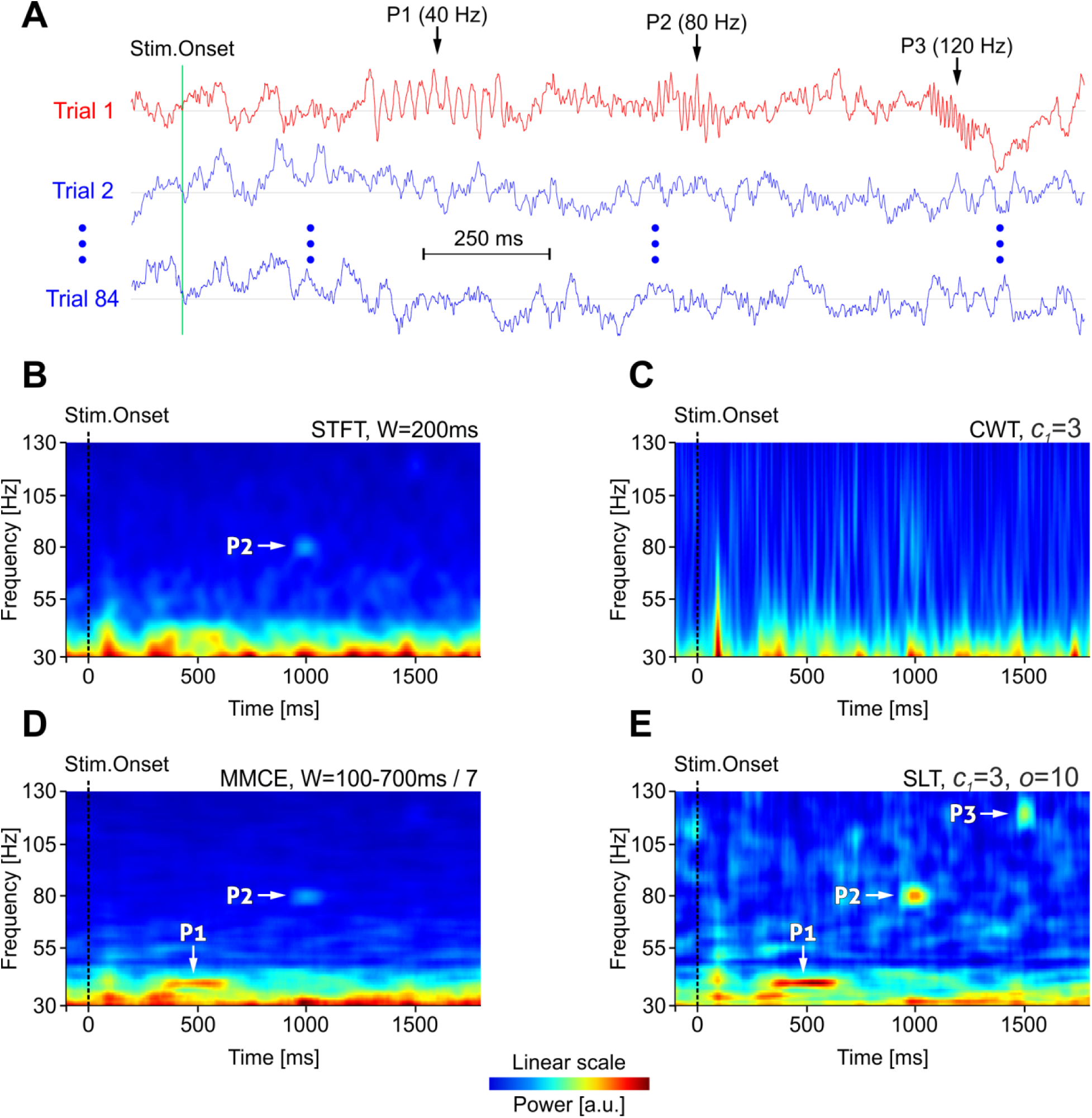
Detection of single-trial gamma bursts. (**A**) Three target packets extending 11 cycles and with frequency 40Hz (P1), 80Hz (P2), and 120Hz (P3) were inserted into a single trial of a set of 84 trials of EEG recordings (Pz electrode). Real data shown. Averaged time-frequency spectrum over the 84 trials using: (**B**) STFT, (**C**) CWT, (**D**) MMCE, (**E**), multiplicative SLT. Arrows show the time-frequency location of detected target packets.

## Discussion

Increasing the resolution of joint time-frequency estimation, especially for the case of non-stationary signals has been a very active field of research in the past decades ^19^. Notable techniques include the Fourier-based MMCE ^17^ and its Probabilistic Latent Component Analysis (PCLA) derivative ^18^. *Superlets* extend these efforts by using a simple, yet powerful wavelet-based approach. They provide remarkable time-frequency resolution by taking advantage of multiple estimates at a range of temporal resolutions and frequency bandwidths. These estimates are combined geometrically (optimal entropic criterion) to evaluate the temporal and frequency location of finite oscillation packets.

In optics, the term super-resolution refers to the ability to resolve details beyond the diffraction (Rayleigh) limit ^20^ by taking advantage of multiple measurements ^21^. In time-frequency analysis, super-resolution refers to the ability to resolve the joint time-frequency density better than it is possible with a single measurement ^19^. *Superlets* provide super-resolution in the time-frequency sense. Indeed, for a finite oscillation packet, their frequency resolution approaches (but does not go beyond) the theoretical Rayleigh frequency as the order of the *superlet* is increased (see Fig. 2C). In addition, each single wavelet estimate obeys the Heisenberg-Gabor uncertainty principle but, since the digitized signal is stored, multiple evaluations can be performed and combined to transcend the time-frequency resolution of each individual estimate. For only frequency super-resolution other techniques are applicable, e.g., based on model fitting ^32^, polyphase analysis filter banks ^33^, Pisarenko harmonic decomposition^34^, or multiple signal classification (MUSIC) ^35^. However, these frequency super-resolution techniques ignore the temporal component and focus on the frequency dimension only.

*Superlets* use the geometric mean (GM) across a set of wavelet responses to sharpen time-frequency localization. Intuitively, the GM “correlates” responses with high temporal precision with those with high frequency precision ^16^. For example, if a large bandwidth wavelet (many cycles) detects a narrow frequency component this will be vetoed out in time if the narrow wavelet at a certain location has a low response, and vice versa. Quantitatively, it has been shown that using the geometric mean to combine individual measurements improves the estimate of the joint time-frequency density and is optimal in a cross-entropy sense ^17,22^. This property is not shared by the arithmetic mean (see Fig. S3), which corresponds to the minimum mean-squared solution ^17^.

The frequency resolution limit for a finite oscillation packet depends on the packet’s duration but temporal resolution can be increased by increasing sampling rate. Typically, LFPs are obtained by low-pass filtering (@300Hz) the electrophysiology signal sampled at much higher rates (32-50kHz) and then downsampling the signal. When using *superlets*, one should keep a high sampling rate after downsampling (e.g., 2-4 kHz) to enable the method to resolve very fine time-frequency details (see Fig. 5B, Fig. 6, and Fig. 8).

Cortical responses exhibit a significant trial-to-trial variability ^36^. Therefore, results are typically averaged across multiple trials. For time-frequency analysis this can pose significant problems ^27,37,38^, for several reasons. First, perceptual processes may be supported by high-frequency gamma bursts whose expression is not necessarily locked to the external events available for aligning the analysis (stimulus onset, button press etc.). As a result, gamma packets may be scattered throughout the time-frequency spectrum and will not sum up coherently in the average. Second, due to the time-frequency uncertainty, isolated packets can be masked out by strong neighboring packets whose estimate leaks over the target’s representation. Because they concentrate the joint time-frequency estimate in each individual trial and in a frequency-specific manner, *superlets* provide a much sharper image of the time-frequency landscape, revealing oscillation packets that remain hidden from other estimation methods (STFT, CWT, MMCE).

Our results indicate that, for averaged time-frequency spectra, traditional methods (STFT, CWT) may fail to reveal the true time-frequency structure within a certain band if strong spectral neighbors exist, as the representation of the latter leaks into the band of interest, compromising its estimation. Powerful oscillation packets can in principle be detected by many methods, albeit with higher precision by *superlets*. However, estimating the surrounding, weaker packets, turns out to be difficult for classical methods, potentially impairing important discoveries about the simultaneous coordination of rhythms across neighboring bands. *Superlets* provide a robust and elegant solution to this problem because the time-frequency concentration of spectral power they provide minimizes the cross-band contamination during spectral estimation.

Compared to other time-frequency superresolution methods, *superlets* provide significantly better results. For example, to estimate the joint time-frequency power density, the closest relative of *superlets*, the MMCE, uses a set of multiple windows with variable size. However, for each window, this size is fixed across all frequencies. This leads the MMCE to suffer from the same problem as the STFT: power of higher frequency local bursts is more diluted than that of low frequencies. *Superlets* borrow the advantage of the wavelet transform by estimating local power density with wavelets whose size is a function of the base cycle duration. Thus, each frequency is treated with “its own size” to reveal the joint time-frequency power density within a window that is a function of frequency. This prevents the dilution phenomenon, specific to the class of estimators based on the Fourier transform.

*Superlets* provide super-resolution in the time-frequency space and may become instrumental in discovering new phenomena in many fields of science. They are conceptually simple, generalizing the CWT. Unlike in the STFT or MMCE, choosing the parameters for *superlets* (*c*_1_, *o*) is relatively easy because wavelets adapt to the timescale of each frequency. As we have shown, *superlets* yield better results than other established time-frequency super-resolution methods. Therefore, they may find multiple applications in the analysis of signals whose time-frequency landscape is complex. In particular, neuroscience will benefit from this new method to identify high frequency oscillation bursts, which are transient and can be hidden by traditional analysis methods.

## Materials and Methods

### Experimental data and ethics

*High-density electroencephalography* (EEG – Biosemi ActiveTwo 128 electrodes) data was recorded @1024 samples/s from healthy human volunteers freely exploring visual stimuli consisting of deformed lattices of dots that represented objects and were presented on a 22” monitor (1680×1050@120fps; distance 1.12m). Subjects had to signal a perceptual decision by pressing one of three buttons congruent with perception (“nothing”, “uncertain”, “seen”). A similar protocol was described elsewhere ^23^. Here, we used data from a single subject, including trials with correct, “seen” responses (63 trials). The protocol was approved by the Local Ethics Committee (approval 1/CE/08.01.2018). Data was collected in accordance with relevant legislation: Directive (EU) 2016/680 and Romanian Law 190/2018.

*In vivo electrophysiology* data was recorded with A32-tet probes (NeuroNexus Technologies Inc) at 32 kSamples/s (Multi Channel Systems MCS GmbH) from primary visual cortex of anesthetized C57/Bl6 mice receiving monocular visual stimulation (1440×900@60fps; distance 10cm) with full-field drifting gratings (0.11 cycles/deg; 1.75 cycles/s; contrast 25-100%; 8 directions in steps of 45°, each shown 10 times). Anesthesia was induced and maintained with a mixture of O_2_ and isoflurane (1.2%) and was constantly monitored based on heart and respiration rates and testing the pedal reflex. Within a stereotaxic device (Stoelting) a craniotomy (1×1mm) was performed over visual cortex. To minimize animal use, multiple datasets were recorded over 6-8 hours from each animal. Experiments were approved by the Local Ethics Committee (3/CE/02.11.2018) and the National Veterinary Authority (ANSVSA; 147/04.12.2018). Local field potentials were obtained by low-pass filtering the signals @300Hz and downsampling to 4kHz.

## Acknowledgements

This work was supported by: two grants from the Romanian National Authority for Scientific Research and Innovation, CNCS-UEFISCDI (project numbers PN-III-P4-ID-PCE-2016-0010 and COFUND-NEURON-NMDAR-PSY), a grant by the European Union’s Horizon 2020 research and innovation program – grant agreement no. 668863-SyBil-AA, and a National Science Foundation grant NSF-IOS-1656830 funded by the US Government. The authors wish to thank Wolf Singer and Gal Vishne for insightful comments on the method and manuscript.

## Author contributions

R.C.M. and V.V.M. developed the method. V.V.M. generated the toy data, A.N-D and R.C.M. recorded the test data, H.B. implemented and tested the method. V.V.M., A.N-D., and R.C.M. wrote the manuscript.

## Supplementary figures

**Fig. S1.**
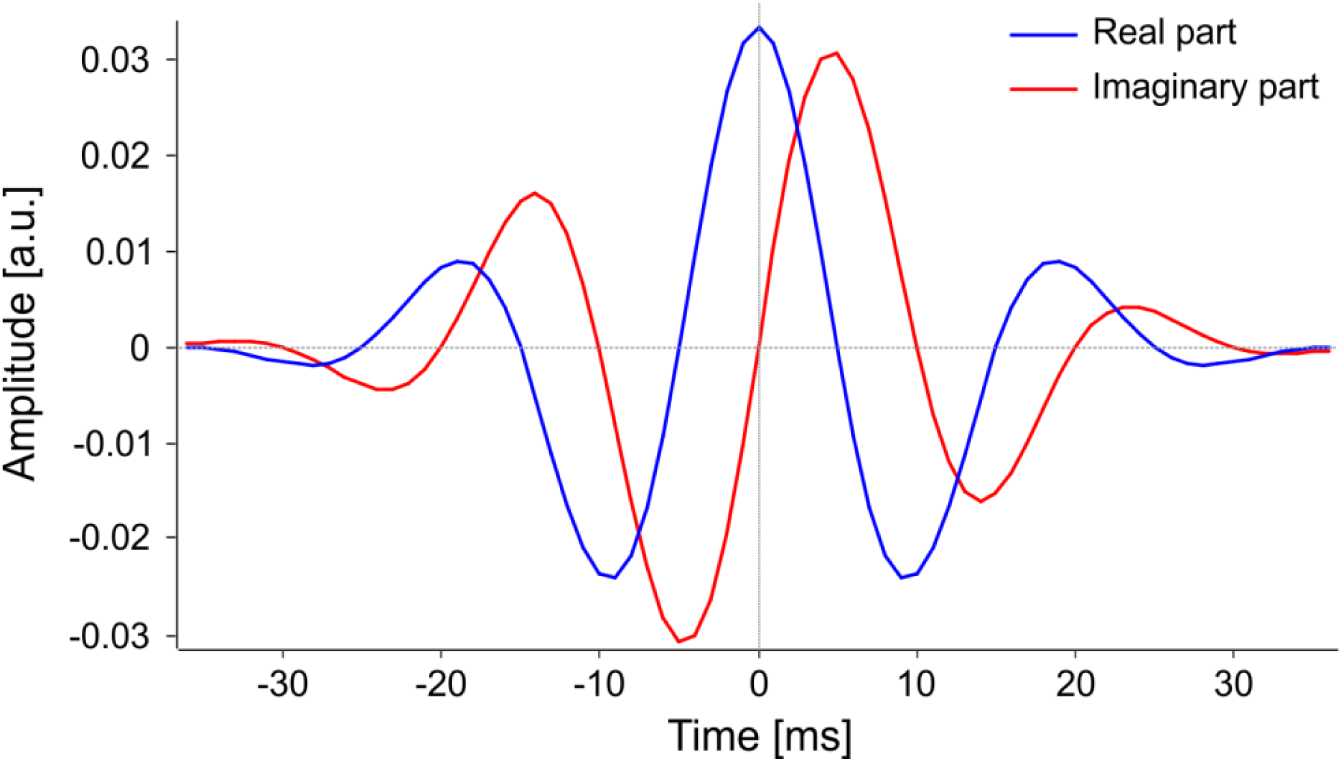
The complex Morlet wavelet. Parameters: *c*=3 cycles and central frequency *f* = 50 Hz, generated at a sampling rate of 1 kHz.

**Fig. S2.**
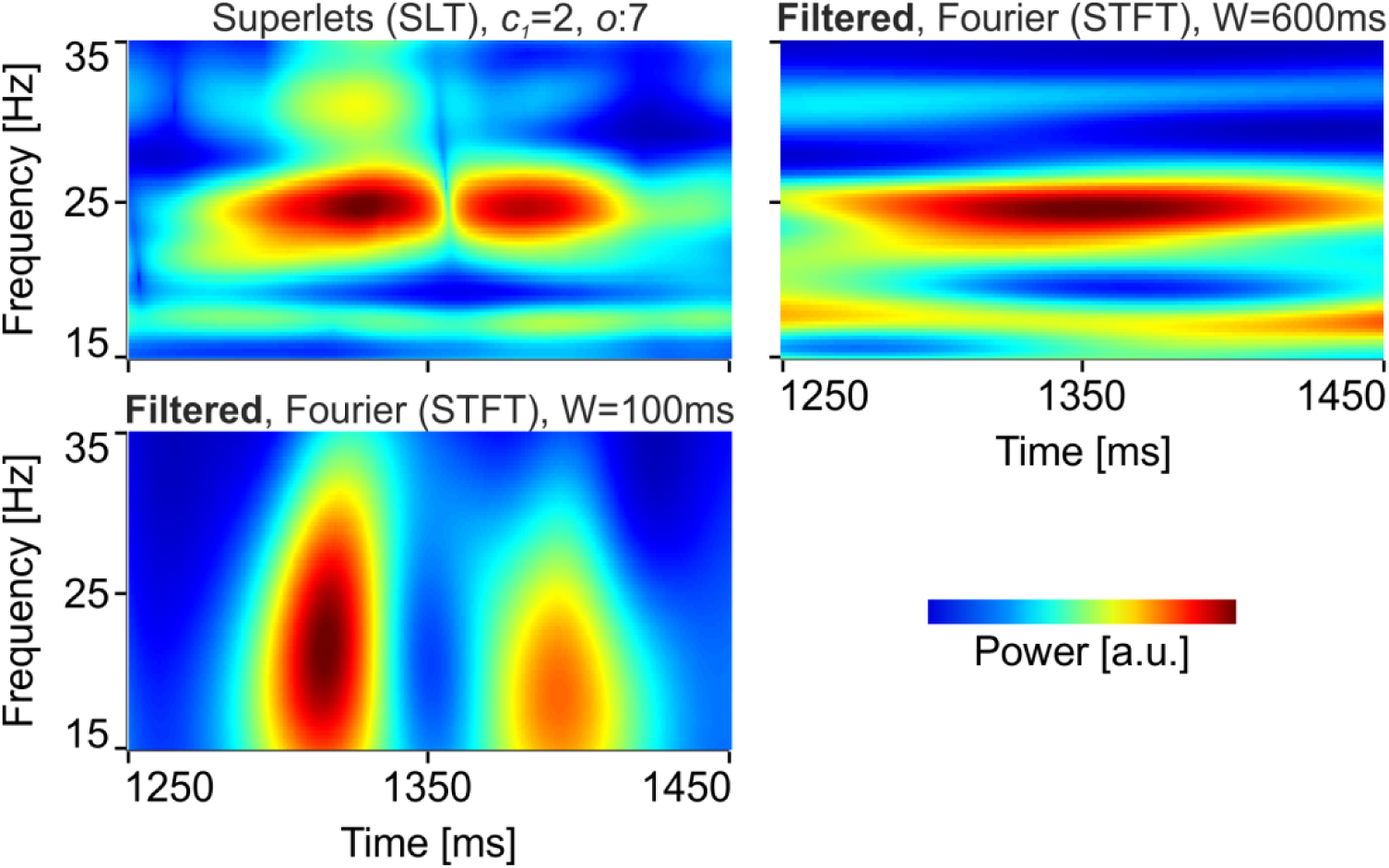
Identification of the oscillation packets revealed by super-resolution using Fourier analysis (STFT). Top-left: super-resolution using multiplicative superlets with *c_1_* = 2, and *o* = 7. Top-right: identification of frequency components using a large Fourier window (W = 600ms). Bottom-left: identification of temporal components using a small Fourier window (W = 100ms). The signal was band-pass filtered at 10-40Hz for the STFT analysis only. See also Fig. 5.

**Fig. S3.**
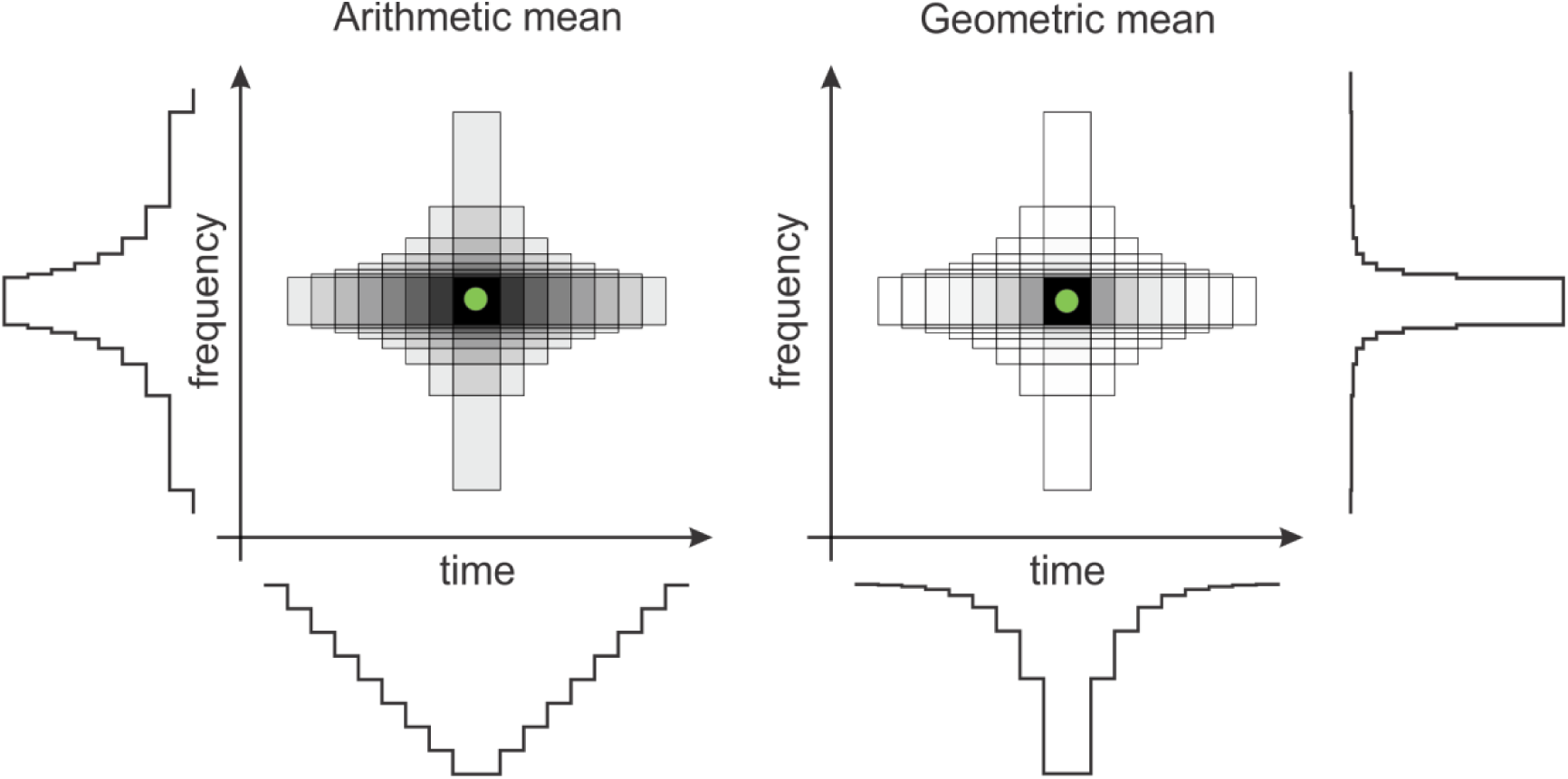
Comparison of arithmetic mean and geometric mean. Estimates using a set of windows with varying frequency and temporal resolution are combined using the arithmetic (left) or geometric (right) mean conveying different time and frequency concentration profiles. In particular, the arithmetic mean suffers from significantly higher temporal smearing than the geometric mean, thus offering a poorer temporal resolution.

